# The ALBA RNA-binding proteins function redundantly to promote growth and flowering in Arabidopsis

**DOI:** 10.1101/758946

**Authors:** Naiqi Wang, Meachery Jalajakumari, Thomas Miller, Mohsen Asadi, Anthony A Millar

## Abstract

RNA-binding proteins (RBPs) are critical regulators of gene expression, but have been poorly studied relative to other classes of gene regulators. Recently, mRNA-interactome capture identified many Arabidopsis RBPs of unknown function, including a family of ALBA domain containing proteins. Arabidopsis has three short-form ALBA homologues (*ALBA1-3*) and three long-form ALBA homologues (*ALBA4-6*), both of which are conserved throughout the plant kingdom. Despite this ancient origin, *ALBA-GUS* translational fusions of *ALBA1, ALBA2, ALBA4*, and *ALBA5* had indistinguishable expression patterns, all being preferentially expressed in young, rapidly dividing tissues. Likewise, all four ALBA proteins had indistinguishable *ALBA-GFP* subcellular localizations in roots, all being preferentially located to the cytoplasm, consistent with being mRNA-binding. Genetic analysis demonstrated redundancy within the long-form *ALBA* family members; in contrast to single *alba* mutants that all appeared wild-type, a triple *alba456* mutant had slower rosette growth and a strong delay in flowering-time. RNA-sequencing found most differentially expressed genes in *alba456* were related to metabolism, not development. Additionally, changes to the *alba456* transcriptome were subtle, suggesting ALBA4-6 participates in a process that does not strongly affect transcriptome composition. Together, our findings demonstrate that ALBA protein function is highly redundant, and is essential for proper growth and flowering in Arabidopsis.

**Highlight:** The RNA-binding ALBA proteins have indistinguishable expression patterns and subcellular localizations in Arabidopsis, acting redundantly to promote growth and flowering via a mechanism that does not strongly affect transcriptome composition.

## Introduction

Post-transcriptional gene regulation is primarily orchestrated via RNA-binding proteins (RBPs). This class of regulators are known to mediate RNA processing, modification, localization, stability and translation/expression (Hentze et al., 2018). Consistent with these fundamental processes, plants contain many hundreds of genes encoding RBPs, being similar in number to genes encoding transcription factors (Silverman et al., 2013). However, very little is known about the molecular and functional roles for the majority of plant RBPs (Cho et al., 2019), with most of our knowledge being derived from bioinformatic extrapolation from other kingdoms (Silverman et al., 2013).

Recently, the mRNA-binding proteome of Arabidopsis was experimentally determined using mRNA-interactome capture, a method using UV-cross-linking of proteins to mRNA, followed by oligo-dT capture of RNA-protein complexes, and then mass spectrometry to identify and quantitate captured proteins (Reichel et al., 2016; Zhang et al., 2016; Marondedze et al., 2016). For example, interactome capture on Arabidopsis etiolated seedlings identified 700 proteins as RBPs, 300 of which were enriched with high confidence. Eighty percent of these proteins contained a bioinformatically predicted RNA-binding domain (RBD), and for the majority this was the first experimental evidence identifying them as RBPs in plants. Amongst these captured RBPs was a family of proteins that contain an “acetylation lowers binding affinity” (ALBA) domain (Bell et al., 2002), where five of the six members were identified as RBPs, four being amongst the most confidently captured proteins (Reichel et al., 2016).

ALBA proteins are highly conserved, being found in all kingdoms of life (Verma et al., 2014). Structures for ALBA proteins have been experimentally determined in the archaea (*Sulfolobus solfataricus*), plant (*Arabidopsis thaliana*) and animal (*Homo sapiens*) kingdoms (Wardleworth et al., 2002, Madej et al., 2014, Wu et al., 2018). All these proteins have similar structures, having dimeric and tetrameric forms which can interact with DNA and RNA (Tanaka et al., 2012, Guo et al., 2014, da Costa et al., 2017, Chan et al., 2018), indicating a conserved nucleic acid-binding capacity. Despite this conserved biochemistry, ALBA proteins in the different kingdoms appear to have diverse molecular functions. In archaea, ALBA proteins ubiquitously bind to DNA, helping to form highly condensed DNA structures, potentially performing analogous functions as eukaryotic histones (Bell et al., 2002, Wardleworth et al., 2002, Laurens et al., 2012, Jelinska et al., 2005). Archaea ALBA proteins also bind RNA, often interacting specifically with double-strand RNA structures, regulating RNA stability (Guo et al., 2003, Guo et al., 2014, Goyal et al., 2016). In protozoa, most studies of ALBA proteins have focused on their RNA-binding capacity (Subota et al., 2011, Chene et al., 2012). For instance, in the trypanosome Leishmania, ALBA proteins can specifically regulate the stability of *AMASTIN* which encodes a transmembrane glycoprotein in a particular development stage, contributing to the control of developmental stage and asexual reproduction (Dupe et al., 2014, Perez-Diaz et al., 2017). In animals, ALBA proteins specifically interact with tRNA. Two ALBA family members, named Rpp20 and Rpp25, are subunits of the Ribonuclease P (RNase P) holoenzyme. Here, an Rpp20/Rpp25 heterodimer specifically bind with pre-tRNA, which is required for the tRNA maturation process performed by the RNase P complex (Hands-Taylor et al., 2010, Reiner et al., 2011).

In plants, little is known regarding the ALBA proteins family. For Arabidopsis, there are six *ALBA* genes, three of which have a long-form structure that consists of an N-terminal ALBA domain, followed by a C-terminal region that possesses multiple Arginine-Glycine (RGG) repeats (At1g76010, At1g20220, At3g07030). The RGG repeats are RNA recognition motifs that specifically interact with guanine (G)-quadruplexes on RNA (Vasilyev et al., 2015, Ozdilek et al., 2017). The other three ALBA proteins are short-forms, and almost solely consist of the ALBA domain (At1g29250, At2g34160, At3g04620). Such long- and short-form structures are found in other kingdoms, such as the protozoan *Trypanosome brucei* (Subota et al., 2011), suggesting an ancient origin of these different ALBA forms.

Currently, information regarding plant ALBA function is only just emerging. Firstly, T-DNA insertional mutations in ALBA long-form genes of either Arabidopsis (At1g76010 and At1g20220) or the liverwort *Marchantia polymorpha*, inhibited root hair growth (Honkanen et al., 2016). In another study, the short-form ALBA proteins were found to bind DNA-RNA hybrids *in vitro*, that they localized to both the nucleus and cytoplasm, where they could form either homo- or heterodimers with one another (Yuan et al., 2019). In the nucleus, they were shown to be R-loop readers, binding throughout the genome at locations consistent with the presence of R-loops, where their function is to stabilize the genome (Yuan et al., 2019). Therefore, the short-form ALBA proteins appear to have a clear role in the nucleus related to DNA-binding. This includes a nuclear localized ALBA protein in rice, whose expression is upregulated by water-deficient conditions (Verma et al., 2014). Other rice *ALBA* homologs were also upregulated under stress conditions or treatment with hormones, suggesting a role in stress-adaptation and other physiological processes (Verma et al., 2018). However, plant ALBA proteins must also have an RNA-binding role. They were amongst the most enrich proteins in mRNA-interactome capture (Reichel et al., 2016), and in an RNA-affinity purification experiment, an ALBA protein (At1g76010) was amongst the most enriched proteins (Gosai et al., 2015).

Here, we perform an initial molecular and functional analysis of the members of the Arabidopsis ALBA family. We find that despite the short- and long-form ALBA proteins having an ancient origin, they have highly similar expression patterns. Analysis of *alba* T-DNA mutant, shows extensive genetic redundancy exists between the different ALBA proteins, where they appear essential for rosette growth and flowering-time in Arabidopsis. Surprisingly, mutation of an entire ALBA clade did not strongly affect transcriptome composition.

## Materials and methods

### Plant material and growth

The wild-type *Arabidopsis thaliana* involved in this project is the Columbia-0 ecotype (Col-0). The *alba4* (SALK_015940), *alba5* (SALK_088909) and *alba6* (SALK_048337) single mutants were obtained from the Arabidopsis Biological Resource Center (ARBC). The seeds were sterilized by chlorine gas for four hours in a sealed desiccator jar and then were germinated on 1/2 Murashige and Skoog (1/2 MS) medium with 7 g/L agar. Growth conditions were either a long-day (16 hours light) or short-day (10 hours light) photoperiod at approximately 100 μmol photons s^-1^ m^-2^ at 22°C. After approximately 10 days, seedlings were transplanted to Debco® plugger soil with Osmocote® Extra Mini fertilizer (3.5 g/L soil) and Azamax® pesticide (0.75 mL/L soil).

### RNA extraction, qRT-PCR and RNA sequencing

Samples were processed in 1.5 mL centrifuge tubes being immediately frozen with liquid nitrogen, followed by being finely grounded with plastic pestles. Total RNA was extracted using TRIzol® (1 mL per 500 mg sample). 14 μg RNA was treated with 14 μL of RQ1 RNase-Free DNase (Promega) and 1 μL of RNaseOut^™^ Recombinant RNase Inhibitor (Invitrogen) following the manufacturer’s protocol. The RNA was then purified using the QiAgen RNeasy mini kit following the RNeasy column clean-up protocol. The RNA quantity and quality were determined via NanoDrop and agarose gel electrophoresis. cDNA was prepared using SuperScript® III Reverse Transcriptase (Invitrogen), with the addition of RNaseOut^™^. For qRT-PCR, 0.4 μL 10 μM specific primer pairs (mixture of forward and reverse primers) was mixed with 10 μL SensiFAST SYBR (Bioline) mastermix and 9.6 μL of cDNA. All the qRT-PCR reactions were performed in three technical replicates, carried out by a QIAGEN Rotor-Gene-Q real-time PCR machine. The comparative quantity of each sample was analyzed with the Rotor-Gene 6000 series software (QIAGEN). The *CYCLOPHILIN* (At2g29960) was used to normalize the mRNA quantities.

For RNA-seq, total RNA was extracted from three biological replicates of seven-day-old *alba456* and *Col-0* seedlings. Quality of purified RNA samples was determined by Nanodrop, agarose gel electrophoresis and via LabChIP GXII (PerkinElmer) analysis. Library preparation, RNA sequencing, differentially expressed gene analysis and GO enrichment analysis were all performed by Beijing Genomics Institute (BGI), Hong Kong.

### Generation of *ALBA:GUS* or *ALBA:GFP* transgenic *Arabidopsis*

Primers were designed amplify genomic fragments of ALBA genes encompassing 5’ regions, exons/introns to the end of the gene except the stop codon (Table S2). *attB1* and *attB2* sites were included to enable Gateway cloning. The amplification of *ALBA* genome sequences was carried out with high-fidelity KOD Hot Start DNA Polymerase (Merck Millipore), according to the manufacturer’s protocol. Amplicons of correct size were gel purified with Wizard® SV Gel and PCR Clean-Up System (Promega). Amplicons were cloned into *pDONR/Zeo* (Invitrogen) using the Gateway BP Clonase II enzyme mix (Invitrogen), and transformed into *Escherichia coli* α-select chemically competent cells (Bioline) via heat shock. Plasmids were screen via restriction enzyme analysis and then entire insert was sequenced to ensure no amplification errors. All correct *ALBA* sequences were then subcloned into *pMDC111* and *pMDC164* destination vectors (Curtis and Grossniklaus, 2003) separately to generate expression clones using Gateway LR Clonase II enzyme mix (Invitrogen). The ALBA-reporter gene junction was verified via DNA sequencing to ensure the fusion gene was in frame. Expression clones were transformed into *Agrobacterium tumefaciens GV3101* via electroporation and were selected on LB plates containing Rifamycin (50 μg/mL), Gentamicin (25 μg/mL) and Kanamycin (50 μg/mL) and verified by restriction enzyme digestion and used to transform Arabidopsis by standard procedures (Clough and Bent, 1998).

### GUS staining

The *ALBA-GUS* transgenic *Arabidopsis* seedlings were fixed with cold 90% acetone for 20 minutes and washed three times with 1X PBS. Then they were vacuum infiltrated in X-Gluc solution [1 mg/mL X-Gluc, 1.66% N,N-dimethyl formamide, 2% ferricyanide (5 mM), 2% ferrocyanide (5 mM), 50% Triton X-100 (0.3%), 4% phosphate buffer (0.5 M), 20% methanol], and incubation at 37°C for 2 hours. Seedlings were then de-stained with 70%-80% ethanol and observed and photographed using a Leica M205C fluorescence microscope.

### DAPI staining and visualization of GFP

The *ALBA-GFP* transgenic *Arabidopsis* seedlings were placed on slides, fixed in 0.1% triton X-100 (diluted in PBS) for 15 minutes and then washed three times in 1X PBS. Samples were then stained by 1 μL/mL DAPI (4’,6-diamidino-2-phenylindole) stored in the dark until confocal microscopy. Visualization and photography were performed using the Zeiss LSM780 or the Leica SP8 confocal microscope. The GFP was excited under 488 nm laser and was observed under 500 ∼ 530 nm spectral detection. The DAPI was excited with 405 nm laser and was observed under 460 spectral detection

### Genotyping and phenotyping of T-DNA Arabidopsis mutants

DNA was extracted from *Arabidopsis* young rosette leaves (Edwards et al., 1991), and the presence of T-DNA alleles or wild-type alleles was tested via PCR using gene-specific and the LBb1.3 primers (Table S3) using the GoTaq® Hot Start Polymerase (Promega). The program was: denaturation at 95°C for 2 minutes; followed by 95°C for 45 seconds, 55°C for 45 seconds and 72°C for 90 seconds for 35 cycles; then the extension 72°C for 5 minutes. The PCR products were analyzed on 1% agarose gel and the position of T-DNA insertions were verified via sequencing the DNA amplicons.

For phenotyping, mutant and wild-type (Col-0) plants were planted side-by-side on the same tray and seed was collected from these plants. These seeds were then sown on 0.5X MS-agar plates. For determining the germination rate, seeds were assessed every 15 hours after being placed in the growth chamber, and germination was defined as radicle protrusion from the seed. At 8-9 days old, seedling were then were transplanted side-by-side into trays of soil comprising of 30 individuals (5 columns X 6 rows). The rosette area was measured via a LemnaTec Scanalyzer every two days at 11 am to 13 pm to ensure the measurements were consistent over the growth period. No further measurements were made once the rosette overlapped with each other on the trays. The flowering-time, the shoot growth and the number of rosette leaves were recorded and counted. The flowering was considered to have occurred when the bolting shoot reached 1 cm (Torti et al., 2012). The rosette leaf were defined as flat leaf with a distinct petiole (Boyes et al., 2001). The counting of rosette leaves was performed every two days at 11 am to 13 pm.

### Alignment of ALBA protein sequences

Amino acid sequences were obtained from Phytozome (Goodstein et al., 2012) using keyword searches for ALBA and then the BLAST Tool was used to verify that there were no other ALBA proteins without annotation. The whole protein sequences of all the ALBA proteins of the species of interest were aligned using three multiple sequence alignment programs Clustal Omega (http://www.ebi.ac.uk/Tools/msa/clustalo/), MUSCLE (http://www.ebi.ac.uk/Tools/msa/muscle/) and COBALT (https://www.ncbi.nlm.nih.gov/tools/cobalt/re_cobalt.cgi). The FASTA format outputs were opened in BioEdit (version 7.2.5; http://www.mbio.ncsu.edu/BioEdit/bioedit.html) to visualise and compare. This comparison allowed identification of consistently aligned regions and alternative alignments of problematic areas with closely clustered gaps. The best sequence with the least problematic areas and the alignment of the Alba domain with the least gaps was selected. This was trimmed down to the Alba domain using the NCBI Conserved domain search (https://www.ncbi.nlm.nih.gov/Structure/cdd/wrpsb.cgi) to identify the extent of the Alba domain and then the “strip columns containing gaps” function in BioEdit was used. The trimmed alignments were uploaded to IQTREE (http://iqtree.cibiv.univie.ac.at/). The job was run at default settings for protein sequences. Trees were visualised using FigTree (v 1.4.3; http://tree.bio.ed.ac.uk/software/figtree/).

### Statistical and bioinformatics analysis

For qRT-PCR, rosette area size, and rosette leaf number, one-way analysis of variance (ANOVA) was carried out to test whether the traits differed significantly between samples or genotypes. Additionally, before ANOVA analysis, the developmental curves of rosette area size or rosette leaf number were fitted to linear models. After ANOVA, the comparisons between each sample or genotype were performed with Tukey’s honest significant difference (HSD) test. The adjusted p-value (*p*) < 0.05 referred to the statistically significant difference. All analyses and plots were made in R (v3.4.3) and RStudio with the package *ggplot2* v3.1.0 (Wickham, 2016). The linear model was fitted using the *lm* function. The ANOVA and Tuckey’s HSD test were accomplished with the package *multcomp* v1.4-10 (Hothorn et al., 2008). For flowering-time data, the student t-test was performed to compare the different genotypes, using R (v3.4.3) and RStudio with the package *ggpubr* v0.2. Whether segregating populations corresponded to Mendelian ratios was determined by Chi-square test.

## Results

### Two distinct clades of ALBA proteins exist throughout the plant kingdom

In Arabidopsis, there are six ALBA genes (Figure 1). Two previous reports have given them different names (Honkanen et al., 2016; Yuan et al., 2019). Although Honkanen et al. (2016) study was first, given the extensive analysis of Yuan et al. (2019), we will follow their ALBA gene nomenclature. There are three shorter-form *ALBA* genes, *ALBA1* (AT1G29250), *ALBA2* (AT2G34160) and *ALBA3* (AT3G04620); and three longer-form *ALBA* genes being *ALBA4* (AT1G76010), *ALBA5* (AT1G20220) and *ALBA6* (AT3G07030) (Figure 1). Phylogenetic finds these shorter and longer forms fall into two distinct clades (data not shown).

**Figure 1.**
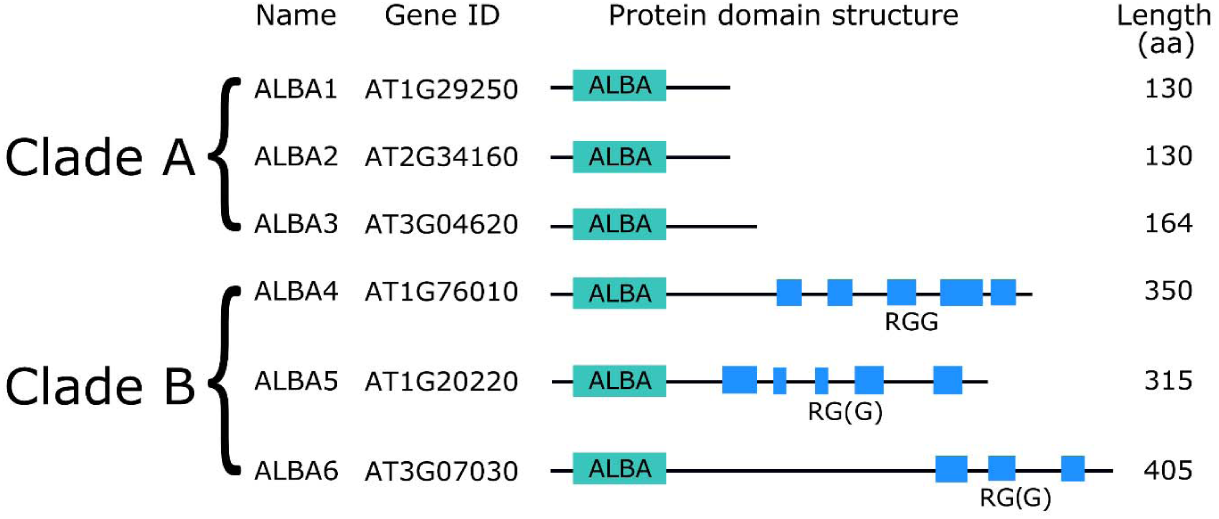
The Arabidopsis ALBA proteins. AT number, domain structure and amino acid length are shown for each gene.

To investigate whether these two clades exists throughout the plant kingdom, ALBA protein sequences were obtained and analysed from dicotyledonous species (*Arabidopsis thaliana, Medicago truncatula, Populus trichocarpa* and *Mimulus guttatus*), monocotyledonous species (*Oryza sativa* and *Sorghum bicolor*), the basal angiosperm *Amborella trichopoda*, the bryophyte *Physcomitrella patens* (moss) and the single celled green alga *Chlamydomonas reinhardtii*. All protein sequences were obtained from Phytozome, aligned using MUSCLE, trimmed to a 68 amino acid alignment of the ALBA domain and used to generate a phylogenetic tree. It clearly shows that these two clades are found throughout the plant kingdom, which we have named Clade A and Clade B (Figure S1).

Clade A proteins are generally shorter with a mean length of approximately 141 amino acids, predominantly consist of a single ALBA domain and have a conserved amino acid sequence (NRIQVS) at the start of their ALBA domains. Clade B proteins are generally longer with a mean length of approximately 290 amino acids and most possess a characteristic domain structure of an ALBA domain followed by a more variable region that contains multiple Arginine-Glycine (RGG) repeats (Figure 1). This clade also has a different conserved amino acid sequence (NEIRIT) at the start of the ALBA domain. The tree implies the two different clades arose before the origin and diversification of plants, suggesting they are ancient and fundamental for plant cellular life. Curiously, in the species we examined, there are equal or near-equal numbers of Clade A and Clade B homologues (Figure S1).

### Transcript expression analysis of the Arabidopsis *ALBA* genes

To initiate a molecular characterization of the *ALBA* gene family in Arabidopsis, ecotype Columbia-0 (Col-0), we used qRT-PCR to quantify the levels of *ALBA* mRNAs. In general, *ALBA1* and *ALBA4* have the highest level in Clade A and Clade B respectively (Figure 2). Analysis in different tissues found *ALBA1, ALBA2, ALBA5* and *ALBA6* had similar mRNA levels in vegetative and reproductive parts of the plant (*p*>0.05, ANOVA), whereas *ALBA4* exhibited higher mRNA levels in rosettes (*p*<0.001, ANOVA, Tukey’s HSD). *ALBA3* had higher mRNA levels in flowers (*p*<0.001, ANOVA, Tukey’s HSD), suggesting potential tissue specificity (Figure 2A). Analysis of *ALBA* mRNA levels during rosette development found a general trend of lower mRNA levels as development progressed (Figure 2B). Although this was clearest for *ALBA4*, for the *ALBA1, ALBA2, ALBA5* and *ALBA6* genes, the oldest tissues consistently contained the lowest *ALBA* mRNA levels (Figure 2B). This suggests these *ALBA* genes are all preferentially transcribed in young tissues. By contrast, *ALBA3* mRNA levels remained consistently low throughout rosette development (*p*>0.5, ANOVA) (Figure 2B).

**Figure 2.**
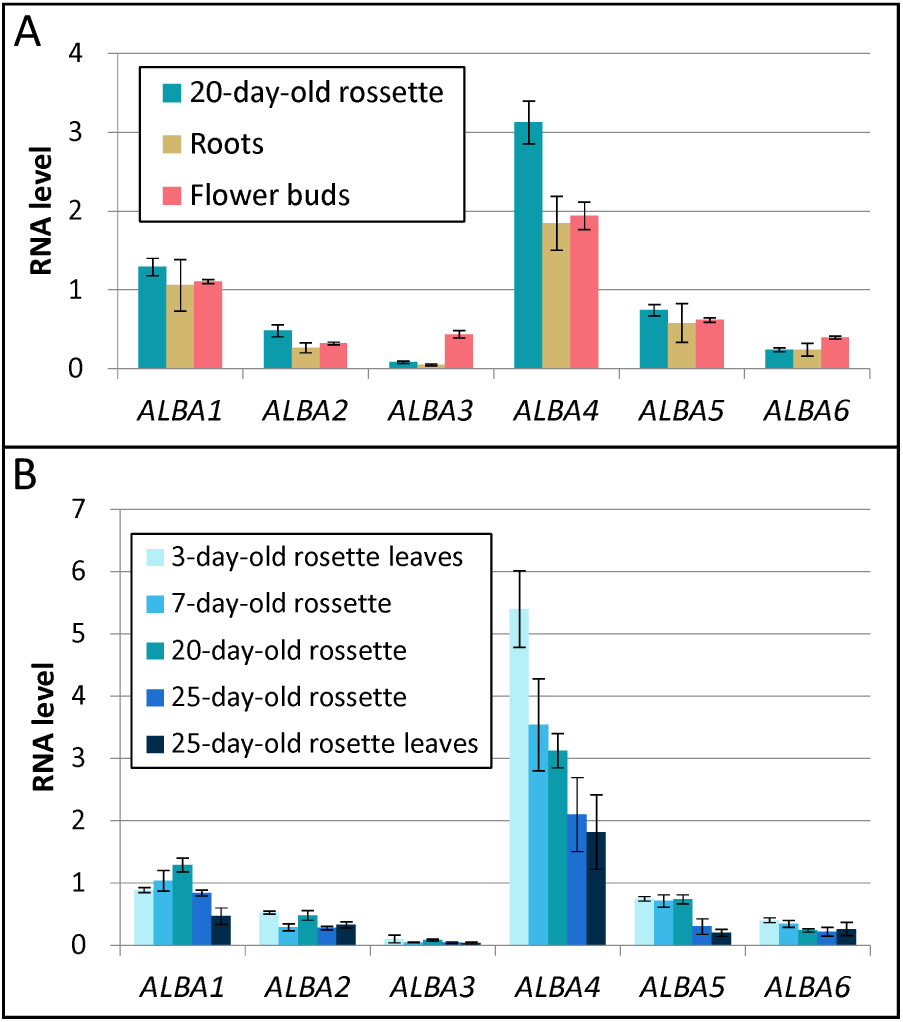
qRT-PCR transcript profiling of the six *ALBA* genes in Arabidopsis. (**A**) mRNA levels in root, rosettes and flowers. (**B**) mRNA levels at different stages of rosette growth. All levels are normalized to *CYCLOPHILIN*. All measurements are the mean of three biological replicates, each of which was determined by three technical replicates (*n*=*3*). The error bars are the standard deviations.

### The Arabidopsis ALBA proteins preferentially express in young tissues

Given that the *ALBA1, ALBA2, ALBA4* and *ALBA5* have the highest transcript levels, protein expression patterns were determined for these four genes. To perform this, these genes were amplified by PCR from Arabidopsis and individually cloned in frame with the *GUS* reporter gene of the pMDC164 vector (Curtis and Grossniklaus, 2003), to generate *ALBA-GUS* translational fusions (Figure 3A). The isolated *ALBA* sequences included the 5’ intergenic region to the preceding upstream gene and all coding region introns (Figure S2, Figure 3A). Including these extensive sequences will help maximize the likelihood that the expression of these *ALBA-GUS* transgenes reflects that of the endogenous *ALBA* genes. Each *ALBA-GUS* transgene was individually transformed into Arabidopsis, as well as an empty pMDC164 vector to act as a negative control.

**Figure 3.**
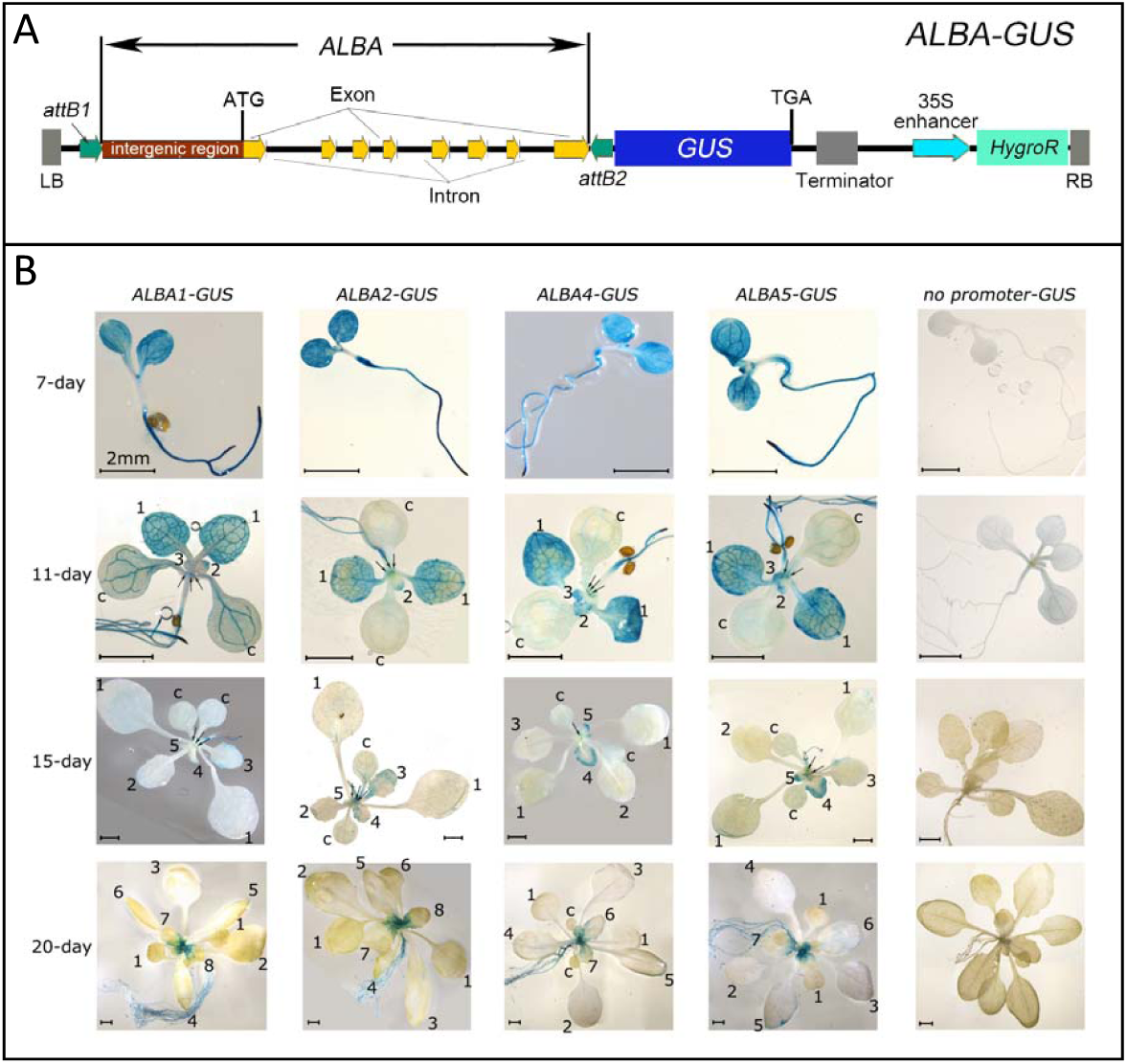
The expression of *ALBA-GUS* transgenes during vegetative development. (**A**) Schematic representation of the *ALBA-GUS* translational fusion for *ALBA1, ALBA2, ALBA4,* and *ALBA5* in the pMDC163 vector. DNA sequences contain 5’ intergenic regions, exon and intron region up, but not including the stop codon, were cloned in frame with the GUS gene. There will be a “scare” of 28 amino acids between the ALBA and GUS. LB = left border, RB = right border, *HygroR* = *Hygromycin* resistance gene. The cartoon is not to scale. (**B**) Expression patterns of *ALBA1-GUS, ALBA2-GUS, ALBA4-GUS, ALBA5-GUS* and *pMDC164* transgenic Arabidopsis throughout vegetative development (7-, 11-, 15- and 20-day old plants are presented). Each picture is representative of at least three independent primary transformants analysed. The order of the leaf emergence is labeled (“c” denotes cotyledon, “1” denotes the first pair of leaves, “2” denotes the second leaf, etc). The vegetative meristem in the shoot apex region is indicated with black arrows. Scale bars = 2 mm.

For each *ALBA-GUS* transgene, GUS staining was performed on multiple independent Arabidopsis transformants that were either 7-, 11-, 15- or 20-days old. In general, all *ALBA-GUS* transgenes had highly similar expression patterns. From 7-to 20-day old plants, GUS activity was consistently present in the shoot apex region (SAR) and the roots, being strongest in the root tips (Figure 3B). Intriguingly, a dynamic expression pattern of ALBA-GUS proteins occurred in leaves. For example, in 7-day-old plants, strong ALBA-GUS expression was found in the cotyledons. However, as the rosette matured, ALBA-GUS expression was lower in mature cotyledons, but was strongly expressed in newly emerging leaves (Figure 3B). Here, ALBA-GUS expression was highest near the leaf margin (Figure S3A), a region that comprises the marginal meristem that controls leaf growth after its emergence (Alvarez et al., 2016). Therefore, consistent with the *ALBA* mRNA levels (Figure 2B), expression appears strongest in young, rapidly dividing tissues. No staining occurred in the negative control plants.

All four *ALBA-GUS* transgenes had highly similar expression pattern in reproductive organs (Figure S3B). In mature flowers, ALBA1-GUS, ALBA2-GUS, ALBA4-GUS and ALBA5-GUS expression patterns appeared indistinguishable from one another in stigmas, filaments, pollen and the veins of sepals. In siliques, ALBA-GUS expression mainly localized to the top and the base of the silique (Figure S3B). Therefore, all four ALBA-GUS expression patterns appeared highly similar, suggesting potential genetic redundancy between these *ALBA* family members.

### ALBA-GFP fusions predominantly localize to the cytoplasm

To investigate subcellular localization, *ALBA-GFP* translation fusions were generated for *ALBA1, ALBA2, ALBA4* and *ALBA5* using the identical gene fragments used to generate the ALBA-GUS fusions, but using pMDC111 as the destination vector (Curtis and Grossniklaus) (Figure 4A). Transgenic Arabidopsis lines were generated for each construct, and expression was observed via con-focal microscopy. An Arabidopsis *35S-GFP* line was used as a control.

**Figure 4.**
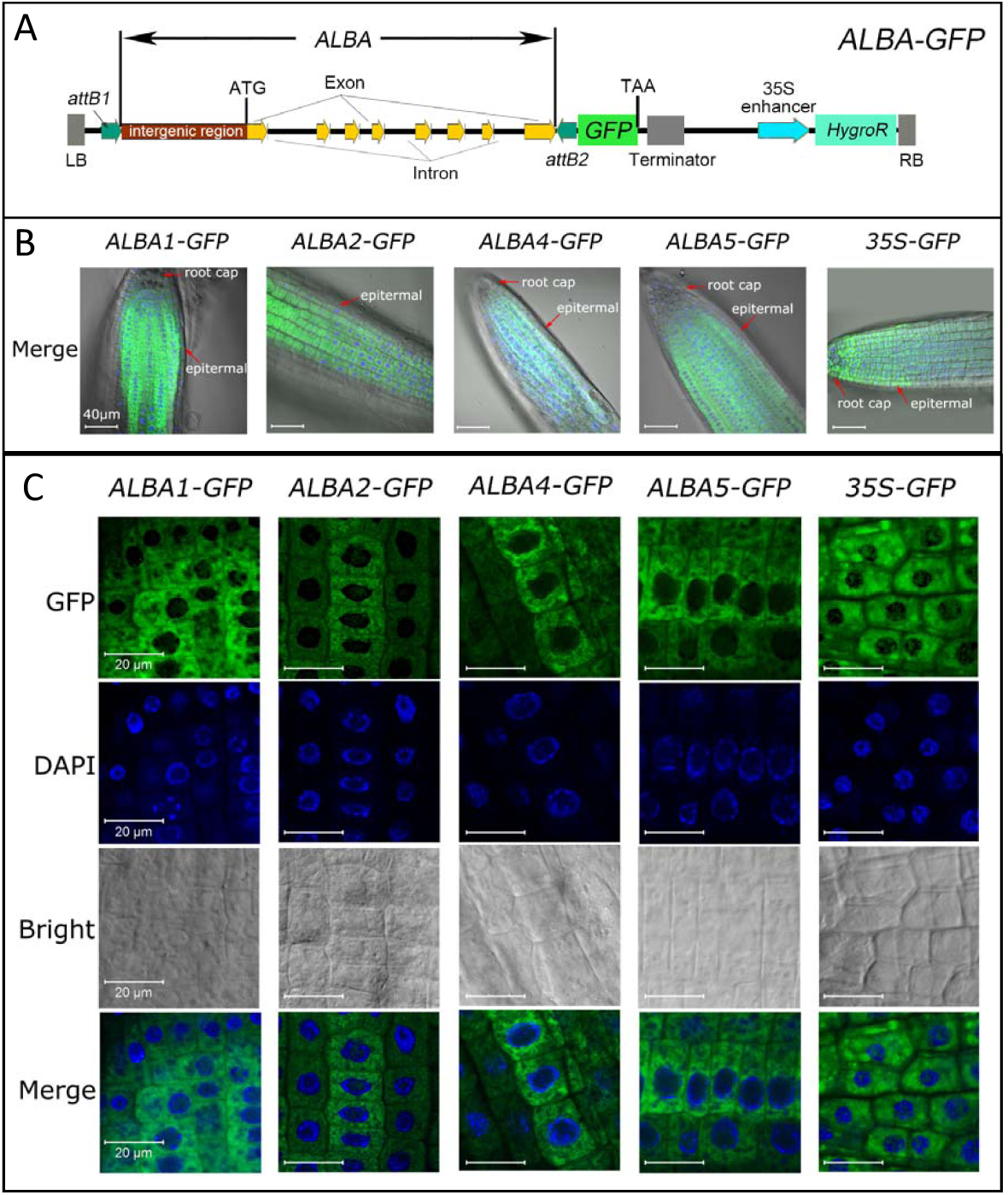
The subcellular localization of the ALBA proteins. (**A**) Schematic representation of the *ALBA-GFP* translational fusion for *ALBA1, ALBA2, ALBA4*, and *ALBA5* in the pMDC111 vector. DNA sequences contain 5’ intergenic regions, exon and intron region up, but not including the stop codon, were cloned in frame with the GUS gene. There will be a “scare” of 24 amino acids between *ALBA* and *GFP*. LB = left boarder, RB = right boarder, HygroR = Hygromycin resistance gene. The cartoon is not to scale. (**B**) *ALBA-GFP* expression in root tips. The GFP fluorescence was green; the nuclei were stained by DAPI, illuminating in blue; the bright field was under transmitted white light. The root cap and the epidermis are indicated with red arrows. Scale bars = 40 μm. (**C**) The GFP fluorescence; DAPI staining and bright field microscopy of *ALBA1-GFP, ALBA2-GFP, ALBA4-GFP, ALBA5-GFP* and *35S-GFP* root tips. Scale bars = 20 μm.

In *ALBA-GFP* seedlings, the strongest and clearest GFP fluorescence was observed in root tips, as this tissue had low auto-florescence. Here, the expression of all four *ALBA-GFP* transgenes appeared indistinguishable, being expressed the strongest in cells within the meristematic zone, but not in the epidermis nor root cap regions. By contrast, expression of the *35S-GFP* transgene occurred in all root tip cells (Figure 4B).

Under increased magnification, the subcellular localization of ALBA-GFP proteins were determined. Firstly, it was found that ALBA-GFP localization appeared mutually exclusive to nuclei, as determined by fluorescence of DAPI staining (Figure 4C). This indicated that the ALBA-GFP proteins were predominantly localized in the cytoplasm. In contrast, the GFP proteins in the *35S-GFP* control were localized in both the nuclei and the cytoplasm (Figure 4C). The predominant cytoplasmic subcellular localization of ALBA-GFP proteins is consistent with a role of binding mature mRNA, rather than that of DNA. Since chloroplasts also contain DNA, ALBA-GFP localization was examined in leaves. It was found that neither ALBA4-GFP nor ALBA5-GFP overlapped with chlorophyll fluorescence (red signal), indicating they are not localized in chloroplasts (Figure S4). Based on this analysis, ALBA-GFP proteins appear predominantly localized to the cytoplasm.

### Generation of an Arabidopsis *alba456* triple mutant

To initiate the functional characterization of the Arabidopsis *ALBA* genes, we chose to focus on *ALBA* genes from Clade B, and investigate whether they are functioning redundantly. To achieve this, the T-DNA insertional mutants *alba4* (SALK_015940), *alba5* (SALK_088909) and *alba6* (SALK_048337) were obtained from the Arabidopsis stock centre (Alonso et al., 2003). The T-DNA insertions were within the coding region for *alba4* and *alba5*, whereas the T-DNA insertion was within the 5’-UTR region for *alba6* (Figure 5). All three single mutants appeared phenotypically indistinguishable from wild-type Arabidopsis (data not shown). Given the high amino acid homology of these ALBA proteins, similar expression patterns and identical sub-cellular localizations, these three *ALBA* genes are potentially functionally redundantly with one another. To investigate this, two *alba4-alba5-alba6* (*alba456*) triple mutant plants were generated. An *alba456-1* isolate was isolated from an *ALBA4/alba4-alba5/alba5-alba6/alba6* parent, and an *alba456-2* isolate from an *alba4/alba4-alba5/alba5-ALBA6/alba6* parent. Having two different isolates will reduce the chances of background mutations segregating with the *alba* mutations in both instances. To confirm the loss-of-function of *ALBA* function in *alba456* mutants, qRT-PCR on *alba456-1, alba456-2* and Col-0 was performed. The mRNA levels of all three *ALBA* genes have been strongly reduced in the *alba456* mutants, indicating this triple mutant corresponds to a strong loss-of-function *alba* mutant (Figure 5B).

**Figure 5.**
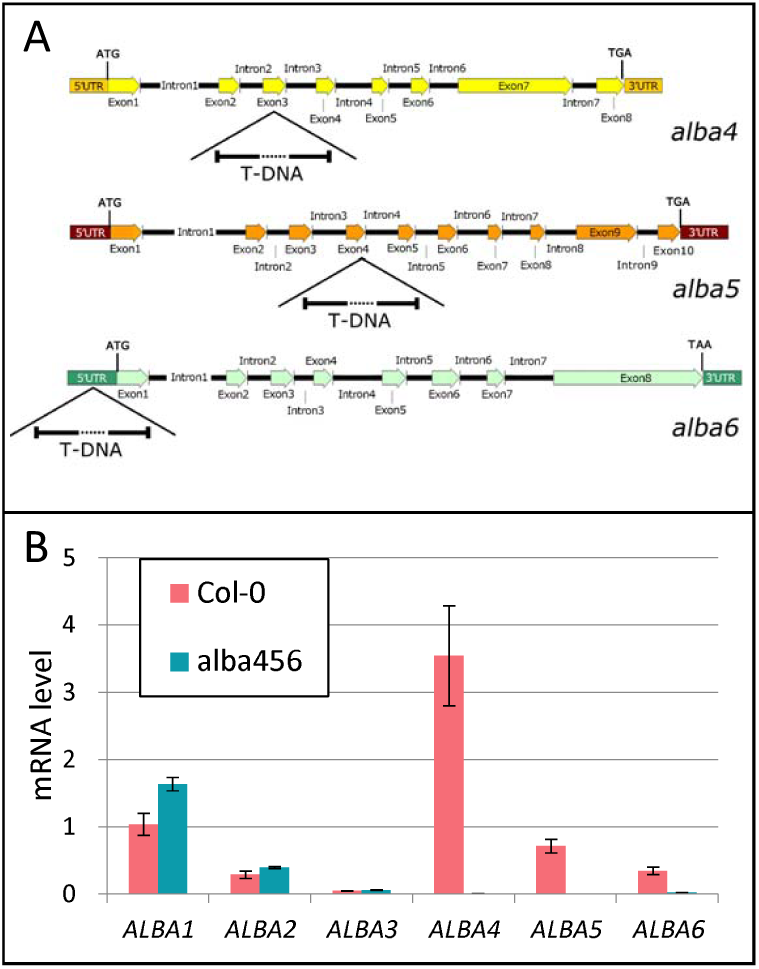
Characterization of the *alba4, alba5* and *alba6* mutants. (**A**) The positions of the T-DNA insertion in the *alba4, alba5*, and *alba6* alleles. The ALBA genes are drawn to scale. (**B**) ALBA mRNA levels x-day old plants of Col-0 and *alba456*. Each measurement represents three biological replicate, with each replicate being composed of three individual plants. RNA levels were normalized to *CYCLOPHILIN.* The error bars represent the standard deviation of the means.

### *alba456* exhibited slower rosette growth and delayed flowering-time

To perform a phenotypic comparison between wild-type (Col-0) and *alba456*, seeds of *alba456-1* and *alba456-2* were sown side-by-side with Col-0 on agar plates. No differences were found in the percentages of seeds that germinated or their germination kinetics (Figure S5). Seedlings were transplanted to soil and the rosette growth of each genotype was monitored by determining the rosette area with a Lemnatech Scanalyser every 48 hours and counting the number of rosette leaves.

From the 16^th^ day to the 26^th^ day, the rosette area of *alba456* grew significantly slower than Col-0 (*p*<0.001, linear mixed model, ANOVA, Tukey’s HSD) (Figure 6A). From the 16^th^ day to the 22^nd^ day, *alba456* had a slightly lower number of leaves than *Col-0* (*p*>0.05, linear mixed model, ANOVA) (Figure 6B), that likely contributes to the smaller rosette area. On average, Col-0 flowered on the 22^nd^ day, whereas *alba456* had an average flowering-time eight days later (*p*<0.01, Student’s t-test) (Figure 6C-D). No significant difference in any of the growth traits was detected between *alba456-1* and *alba456-2* (*p*>0.05, Student’s t-test). Highly similar rosette growth and flowering-time results were obtained in an independent replication (Figure S6). Additionally, similar reductions to rosette growth and delays to flowering-time were seen under short-day conditions, although the differences were not as strong (Figure S7). These experiments argue *ALBA4, ALBA5* and *ALBA6* are required for proper growth and development of Arabidopsis.

**Figure 6.**
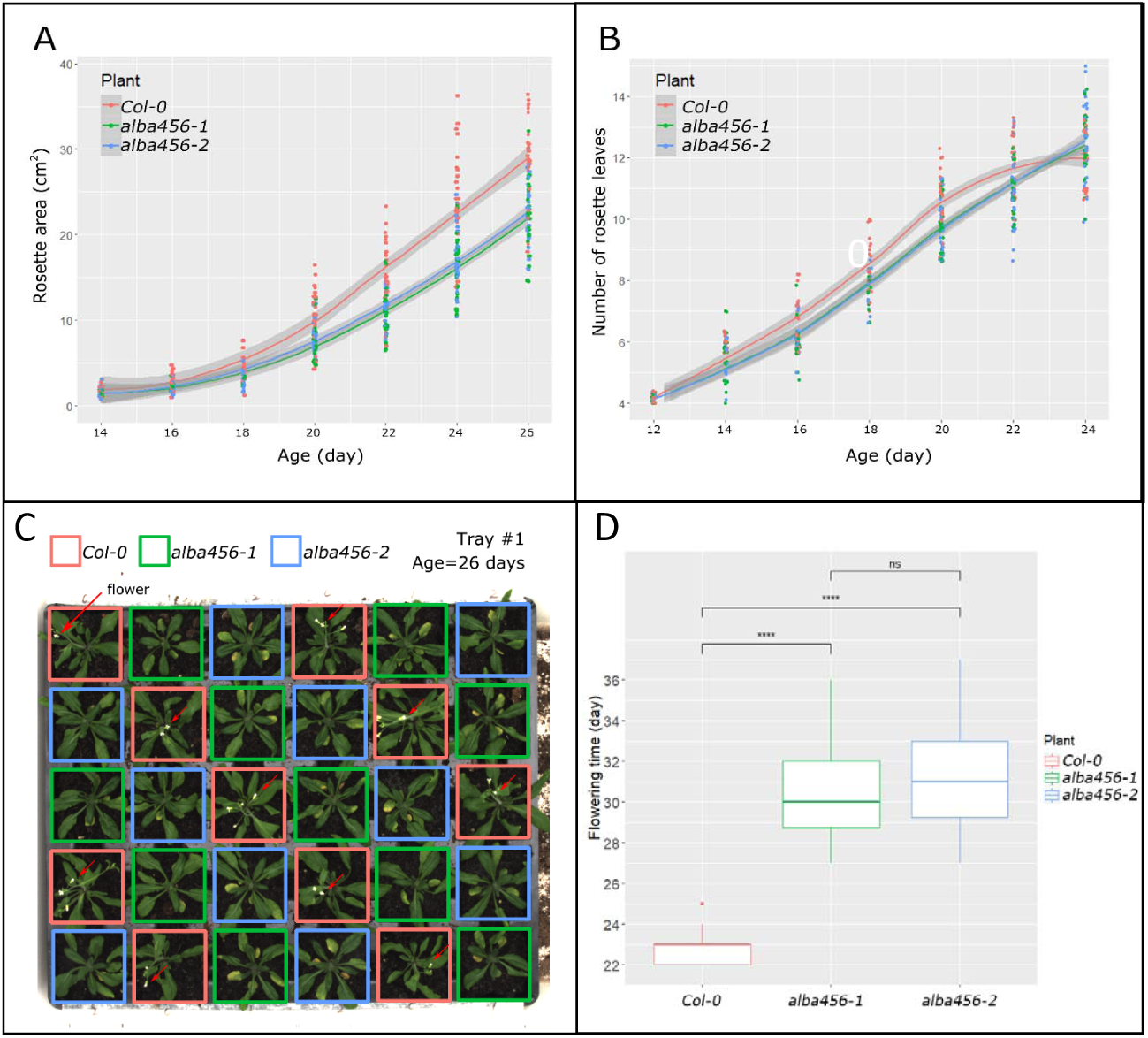
Phenotypic analysis of Col-0, *alba456-1* and *alba456-2*. (**A**) Rosette area from 14-to 26-days old plants. There was a significant difference of the rosette area development between the three groups (*p*<0.001). (**B**) The curves of the rosette leaves number of Col-0 and the mutants. There was no significance between them (p>0.05). For (**A**) and (**B**), the technical replicates are the measurements (*n*=3), the biological replicates are the plants of Col-0 (*n*=28), *alba456-1* (*n*=28) and *alba456-2* (*n*=29). The grey shadow flanking the curve is the confidence interval; the significance of the differences was defined by the ANOVA and Tukey’s HDS following the linear mixed model. (**C**) The flowering-time of Col-0, *alba456-1*, and *alba456-2*. Aerial view of 26-days old plants of the different genotypes. Red arrows indicate flowers. (**D**) The boxplot of the flowering-time. The biological replicates were individual plants in each group (Col-0: *n*=28, *alba456-1*: *n*=28, *alba456-2*: *n*=29). The centerline in the box is the median; the box indicates where the middle 50% of the data lie; the “whiskers” indicate a “reasonable” estimate of the spread of the data. The *alba456-1* and alba456-2 possessed a significantly later flowering-time than Col-0. However, there is no significant difference between *alba456-1* and *alba456-2* (**** denotes *p*<0.0001, “ns” indicates “no significant difference”, analyzed by Student’s t-test).

### The late flower-time phenotype of *alba456* segregated with the *alba* mutations

To determine whether the delayed flowering-time was strictly segregating with the *alba456* mutations, 27 segregating progenies of an *ALBA4/alba4-alba56* and 28 progenies from an *alba45-ALBA6/alba6* mutant were genotyped and scored for their flowering-time (Figure 7). For *ALBA4/alba4-alba56*, five progeny were *alba456* (18.5%), 20 progeny were *ALBA4/alba4-alba56* (74.1%), and two progeny were *ALBA4-alba56* (7.4%). For *alba45-ALBA6/alba6*, one progeny was *alba456* (3.7%), 12 progeny were *alba45-ALBA6/alba6* (42.8%), and 15 progeny were *alba45-ALBA6* (53.6%). Although the segregation did not fit a Mendelian ratio (*p*<0.05, Chi-square test), in both groups the *alba456* progeny had significantly delayed flowering-times (*p<*0.01, analyzed by ANOVA test, Tukey’s HSD). By contrast, plants containing *ALBA4* or *ALBA6* alleles had flowering-times more similar to Col-0 (Figure 7). Thus, this genetically demonstrates that the delayed flowering-time segregated with the *alba* mutations. Furthermore, progeny containing *ALBA4* allele(s) flowered earlier than mutants possessing *ALBA6* allele(s), which suggests *ALBA4* is more predominant than *ALBA6* (Figure 7). This is consistent with the higher mRNA levels of *ALBA4* (Figure 3).

**Figure 7.**
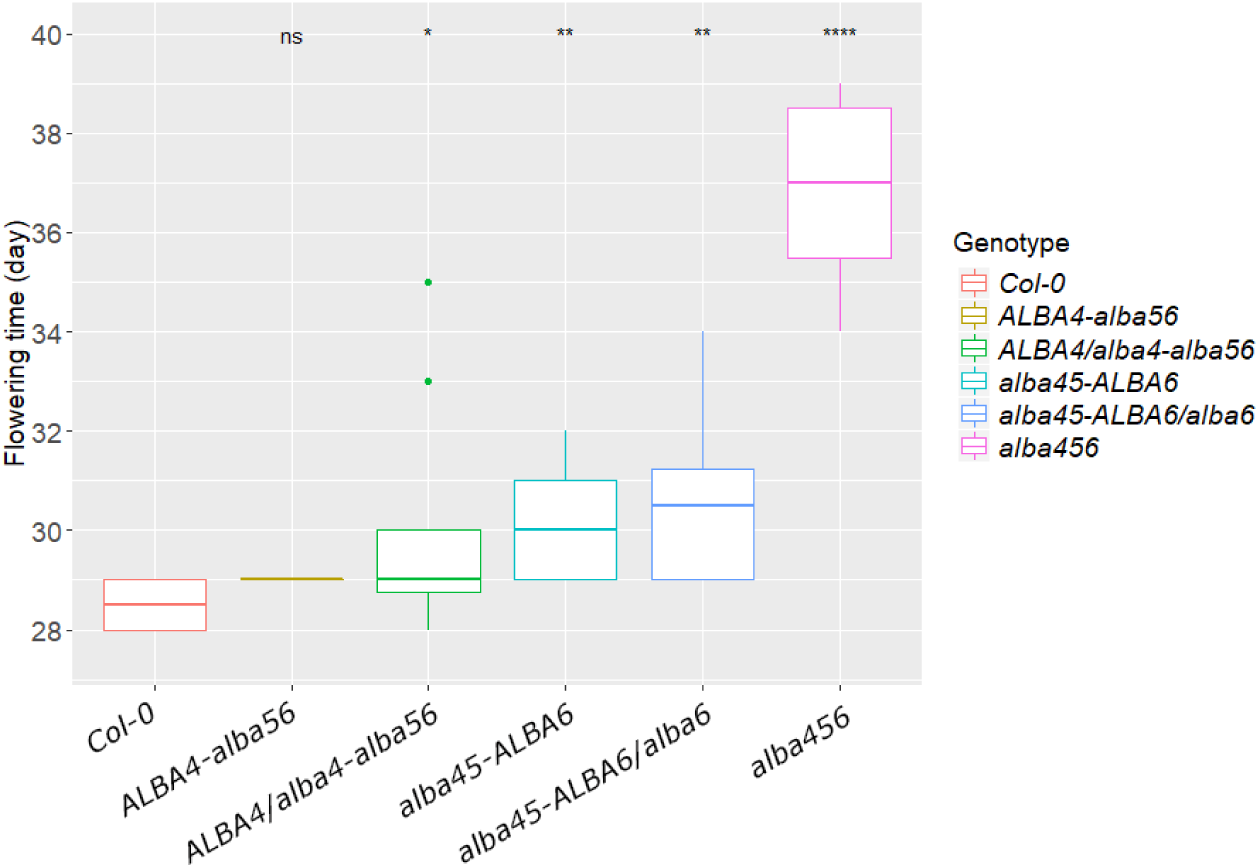
A delayed flowering-time segregates with the *alba456* genotype. Flowering-time were scored from progenies derived from either an *ALBA4/alba4-alba56* (*n*=27) or alba45-*ALBA6/alba6* (*n*=28) parents, as well as Col-0 (*n*=4), all of which were grown under identical conditions. Compared to Col-0, the flowering-time of *ALBA4-alba56* was not significantly different, *ALBA4/alba4-alba56* had slightly delayed flowering (*p*<0.05), *alba45-ALBA6* and *alba45-ALBA6/alba6* exhibited a more significant delayed flowering (*p*<0.01), whereas the alba456 had strongly delayed flowering (*p*<0.0001). The “ns” denotes no significant difference, * denotes *p*<0.05, ** denotes *p*<0.01, **** denotes *p*<0.0001 (Student’s t-test).

### Few mRNAs exhibit high fold-level changes in the *alba456* transcriptome

Since ALBA4, ALBA5 and ALBA6 proteins are experimentally demonstrated mRNA-binding proteins (Reichel et al., 2016), to gain insights into their molecular function, the *alba456* transcriptome was characterized and compared to Col-0. As it was demonstrated that ALBA4 and ALBA5 are strongly and widely expressed in 7-day-old seedlings (Figure 4B), this stage was chosen to perform RNA-seq in a bid to identify the direct effects of the loss-of-function of ALBA4, ALBA5 and ALBA6. RNA from three biological replicates of both Col-0 and *alba456* were prepared and sent to BGI Genomics Co., Ltd for sequencing (BGISEQ-500 platform) and bioinformatic analysis. Over 55 million clean reads were obtained for each of the six samples, for an average of 5.56 Gb bases per sample, with an average mapping ratio of 91.53%, identifying over 23 K gene models.

Firstly, to assess alteration to splicing in *alba456*, differentially spliced genes (DSGs) were identified in the Col-0 and *alba456* transcriptomes. However, only a few genes appeared differentially spliced (Figure S8), indicating that ALBA4-6 were unlikely to play a major or broad role in mRNA processing. Next, differentially expressed genes (DEGs) were identified. Of a total of 23,547 genes, 388 were differentially expressed in *alba456* compared to Col-0 at the two-fold change cutoff, with 173 DEGs being *alba456* upregulated and 215 DEGs being *alba456* downregulated (P < 0.05) (Figure 8). The most downregulated genes in *alba456* were *ALBA4, ALBA5* and *ALBA6* (Table S1), confirming that *alba456* is a strong loss-of-function *alba* mutant. However, the fold-level changes to the majority of the DEGs was modest; at the five-fold change level, there were only six downregulated genes (three ALBA genes, two hypothetical proteins, and a RAS GTP binding protein), and six upregulated genes (Table S1). The identity of these 12 DEGs was uninformative regarding the understanding the *alba456* phenotype.

**Figure 8.**
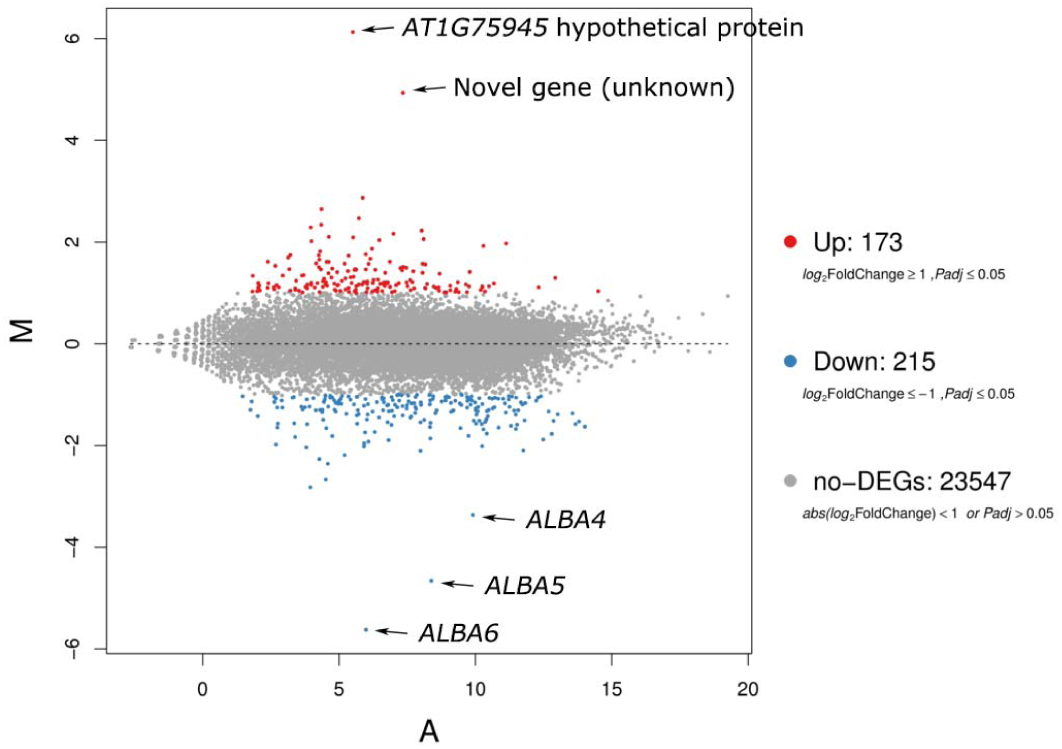
The MA plot of all identified genes in Col-0 and *alba456*. The x-axis represents value A (log2 transformed mean expression level). The y-axis represents value M (log2 transformed fold change in *alba456* compared to Col-0). Red dots represent up-regulated DEGs (M≥1). Blue dots represent down-regulated DEGs (M≤1). Gray points represent non-DEGs.

The DEGs were functionally annotated by Gene Ontology (GO) enrichment analysis. However, for biological process ontology, there was no significant bias in the annotated terms of the DEGs (Figure S9A). For a more detailed analysis, the identity of the top 50 down- and up-regulated genes in *alba456* was determined (Table S1). Some closely related genes appeared co-regulated. For down-regulated DEGs, this includes two highly related members of the LA RELATED PROTEIN family of RBPs, *LARP6a* and *LARP6b*. Also three members of the ROXY family (also known as *GRX5, GRX8* and *GRX11*), are all downregulated. They are involved in cell redox homeostasis, and are all induced by nitrate (Patterson et al., 2016). Conversely, three members of the *FE-UPTAKE-INDUCING PEPTIDE* family, *IRONMAN 1, IRONMAN 2* and *IRONMAN 3*, were upregulated. The expression of these peptide sequences (∼ 50 amino acids) is highly responsive to iron deficiency, where they play a role in iron acquisition and homeostasis (Grillet et al., 2018). Similarly, the *RESPONSE TO LOW SULFUR* (*LSU*) genes, *LSU1* and *LSU3* were both upregulated in *alba456*, and again, both genes encode small proteins (∼95 amino acids) (Lewandowska et al., 2010). Other genes related to low sulfur are also induced, including *SULPHUR DEFICIENCY-INDUCED 1*, and *SULFATE TRANSPORTER 1;3*. Therefore, the expression level changes of all these nitrate, iron and sulfur responsive genes suggest *alba456* plants are experiencing nutrient deficiency. Finally, the *ROXY* genes, and many other DEGs have roles in the oxidation-reduction process (Table S1). Supporting a role of ALBA proteins in this process, the *OsALBA1* gene was shown to play a role in tolerance to oxidative stress, via complementation of a yeast mutant (Verma et al., 2014).

Given this, and the absence of any known developmental genes involved in rosette growth or flowering-time, this data suggested the observed phenotypes is related to alterations to metabolism, rather than developmental programs. This is supported by the classification of DEGs based on the Kyoto Encyclopedia of Genes and Genomes (KEGG) database, which demonstrated that most DEGs corresponded to metabolic pathways (Figure S9B). Considering *alba456* exhibits slower rosette growth, whether these alterations to metabolism are direct or indirect effects from lack of ALBA expression is unknown. However, given the small fold-change levels for the majority of DEGs and the modest numbers of DEGs and DSGs, despite ALBA4-6 being RBPs, their loss does not appear to have a strong and widespread impact on the transcriptome.

## Discussion

### Clade A and Clade B ALBA proteins appear ancient in origin

In this paper, we carry out molecular and functional characterization of the ALBA family of proteins in Arabidopsis. ALBA proteins are found throughout the plant kingdom and separate into two distinct clades; Clade A, which contained short-form ALBA proteins (consists mainly of a single ALBA domain) and Clade B, long-form ALBA proteins (an N-terminal single ALBA domain with a C-terminal region containing multiple RGG repeats). Given the protozoan *Trypanosoma brucei* has analogous short-form and long-form ALBA homologues (Subota et al., 2011), these two ALBA clades must be ancient in origin, and their high conservation implies they are fundamental for cellular life. Curiously, in many different plant species, Clade A and Clade B genes are present in a 1:1 ratio. One possible reason for this is that a short- and long-form ALBA protein specifically dimerize. Although there is evidence of ALBA proteins dimerizing with one another (Yuan et al., 2019), there is no evidence of such specific dimerization.

### The *ALBA* genes are preferentially expressed in rapidly dividing tissues

Despite the apparent ancient origin of the two clades, Clade A (ALBA1 and ALBA2) and Clade B (ALBA4 and ALBA5) proteins have indistinguishable expression patterns (Figure 3), suggesting that these proteins are involved in similar molecular/biological processes. This raised the possibility of functionally redundancy of not only within an ALBA clade, but between Clade A and Clade B members. The *ALBA* genes were expressed highest in young, rapidly dividing tissues. This includes both root and shoot apex regions. For leaves, the expression of all *ALBA:GUS* transgenes exhibited a transient pulse of expression, being highly expressed in newly emerged leaves, but weakly expressed in mature expanded cotyledons/leaves (Figure 3). This leaf expression was highest near the marginal meristem, tissues that promotes leaf distal growth (Figure 3; Figure S3A) (Alvarez et al., 2016). As all these tissues are actively dividing, it would be assumed that they are highly metabolically active, containing high mRNA levels undergoing strong translation which would be required for cellular growth and expansion.

### The ALBA-GFP fusion proteins have similar subcellular localizations

In addition to these highly similar expression patterns, the subcellular localization of ALBA1, ALBA2, ALBA4 and ALBA5 appear identical, all being predominantly located in the cytoplasm when examine in root tips as determined by the expression of C-terminal fusions of GFP to the ALBA proteins. A cytoplasmic subcellular localization is consistent with these proteins being mRNA-binding. However, other reports show conflicting results. One report, using C-terminal ALBA-GFP fusions, found that ALBA1 and ALBA2 were localized to both the nucleus and cytoplasm (Yuan et al., 2019). Another study expressed N- and C-terminal GFP fusions with *ALBA1* and *ALBA2* in Arabidopsis, and in agreement with our analysis found ALBA1 in the cytosol, whereas ALBA2 was either located to the cytosol (C-terminal) or the cytosol and nucleus (N-terminal) (Palm et al., 2016). In both these studies, the ALBA/GFP fusions were transiently expressed with constitutive promoters in mesophyll protoplasts. Similarly, tranisent assay of the rice OsALBA1 protein in epidermal onion cells was located to both the nucleus and cytoplasm (Verma et al., 2014). By contrast, we stably expressed ALBA genes with endogenous promoters and analysed root tips. Such variations in approach could explain the discrepancies between these studies. In other studies using nuclear/cytoplasmic fractionations, ALBA proteins were found in the nucleus as well as the cytoplasm (Yuan et al., 2019, Verma et al., 2014).

In multiple kingdoms, ABLA proteins have been shown to bind both DNA and RNA, and given that Arabidopsis ALBA1 and ALBA2 bind R-loops (Yuan et al., 2019), as well as mRNA (Reichel et al., 2016), it would seem highly likely they are located to both subcellular locations. In other kingdoms, ALBA proteins are found in both subcellular locations. Some ALBA proteins in protozoa are predominantly localized to the cytoplasm (Mani et al., 2011, Chene et al., 2012), whereas some animal ALBA proteins function mainly in the nucleus (Hands-Taylor et al., 2010). In *Leishmania*, some ALBA proteins have a dynamic localization, being located predominantly in the cytoplasm or nucleus depending on the developmental stage (Dupe et al., 2015). Therefore, it is possible that plant ALBA proteins are also differentially localized, which may depend on expression level, developmental stage and/or environmental factors. Further analyses are required to resolve this.

### Perturbed growth and metabolism of the *alba456* mutant

A delayed flowering-time in the *alba456* triple mutant adds to the number of RBPs that have been implicated in controlling flowering-time (Cho et al., 2019, Steffen et al., 2019). As delayed flowering and reduced rosette growth were not apparent in the corresponding single mutants, this demonstrated functional redundancy between *ALBA4, ALBA5* and *ALBA6*. Such an inhibition in growth could be consider consistent with their preferential expression in young, rapidly dividing tissues, where inhibiting the function of these tissues would be expected to negatively impact growth (Figure 2, 3). This is supported by the RNA-seq analysis on the transcriptomes of Col-0 and *alba456*, which found none of the DEGs corresponded to important developmental control genes associated with leaf development or flowering-time (Table S1). This suggests that unlike some RBPs that directly regulate genes in developmental pathways (Steffen et al., 2019), the *alba456* phenotype arises from indirect effects, possibly due to an altered metabolism. Supporting this was the identity of the DEGs and the KEGG enrichment analysis, which found *alba456* DEGs are predominantly related to metabolic pathways, many of which are associated with nutrient deficiency. Therefore, we speculate that perturbation of metabolism slows *alba456* growth, reducing rosette size and delaying flowering-time.

Whether ALBA proteins are directly affecting these DEGs, or whether altered expression of these genes are an indirect consequence of a more general process that is perturbed in *alba456* is unknown. For example, the indirect phenotypic affects were reported for the maize RBP Dek42 that regulates pre-mRNA splicing. The *dek42* mutant predominantly affects starch metabolic processes, but perturbation of this process resulted in seedling lethality (Zuo et al., 2019). The *dek42* mutant caused differential expression to approximately 6% of the transcriptome (Zuo et al., 2019). Similarly, other examples of mutations of RBPs resulted in global changes to the transcriptome. This includes a 14-day old mutant in two RNA recognition motif proteins, RZ-1B and RZ-1C, that had 3,176 DEGs (difference > 1.5-fold, P < 0.01) compared to wild-type (Wu et al., 2016). By contrast, much smaller changes to the *alba456* transcriptome were observed; only 1.6% of the *alba456* transcriptome were DEGs at the 2-fold change cutoff (or 0.05% at the 5-fold change cutoff) and there were very few DSGs (Figure S7). This is despite the *alba456* mutant displaying a clear phenotype, and ALBA4-6 being strongly captured RBPs by mRNA-interactome analysis (Reichel et al., 2016). This may suggest that ALBA4-6 may only be regulating a small cohort of mRNAs. Alternatively, the ALBA proteins may be regulating RNA processes that do not directly affect transcriptome composition, such as translational control (Szostak and Gebauer, 2013). The expression of ALBA proteins in rapidly dividing tissues may support the function associated with translation, as one would assume these tissues would have high levels of translational activity. ALBA proteins from other kingdoms regulate translation. In the protozoa *Leishmania, Trypanosoma* and *Plasmodium*, the ALBA proteins *Li*ALBA20, *Tc*ALBA30 and *Pf*ALBA1 repress the translation of their target mRNAs (Dupe et al., 2014, Perez-Diaz et al., 2017, Chene et al., 2012).

Alternatively, the phenotypes of the *alba456* mutant may not related to their mRNA-binding function, as other ALBA family members have been shown to be associated with R-loops in the nucleus (Yuan et al., 2019), or even possibly playing a role in oxidative stress tolerance (Verma et al., 2014). More work is needed to determine the molecular explanation of the *alba456* phenotype.

## Acknowledgments

We would like to thank Daryl Webb (Centre for Advanced Microscopy) for help with con-focal microscopy, Leila Blackman for help with DAPI staining and Teresa Neeman (Biological Data Science Institute) for the biostatistics advice. We thank the Salk Institute Genomic Analysis Laboratory for providing the sequence-indexed Arabidopsis T-DNA insertion mutants. Funding for the SIGnAL indexed insertion mutant collection was provided by the National Science Foundation. Funds for this project were provided by the Research School of Biology, ANU.

## Supplementary Figure Legends

**Figure S1. Phylogenetic alignment of ALBA proteins from divergent species across the plant kingdom**. *A*. *thaliana* proteins highlighted in green and identified by the code ‘At.ALBA’. The proteins from other species are identified by the code at the start of the gene ID. This includes *Oryza sativa*: ‘LOC_Os’, *Sorghum bicolor*: ‘Sobic.’, *Populus trichocarpa*: ‘Potri.’, *Medicago truncatula*: ‘Medtr’, *Mimulus guttatus*: ‘Migut.’, *Amborella trichopoda*: ‘AmTr_v1.0_scaffold’, *Physcomitrella patens*: ‘Pp’, and *Chlamydomonas reinhardtii*: ‘Cre’. ALBA proteins across the plant kingdom fall into two families with members of both families present in all organisms. Clade A share a characteristic NEIRIT motif at the start of the Alba domain. Clade B shares a characteristic NRIQVS motif at the start of the Alba domain. Node values are ultrafast bootstrap support percentages.

**Figure S2. The amplicons of *attB-ALBA* sequences**. The 5’ intergenic region contained the genomic sequence from the stop codon of the upstream gene to the start codon of the *ALBA* gene. The *ALBA* gene sequences contained all the exons and introns except the stop codon, and no 3’ intergenic sequences were included. The *attB1* and *attB2* sites were incorporated to make the amplicon compatible for Gateway cloning. The lengths of the amplicons are indicated in base pairs.

**Figure S3. GUS staining in leaves and reproductive organs of *ALBA-GUS* transgenic plants. (A)** GUS staining of leaves of *ALBA-GUS* plants. **(B)** GUS staining of mature flowers and siliques of *ALBA-GUS* plants. Each picture is a representative of least three independent primary transformants examined. Scale bars = 1 mm.

**Figure S4. ALBA4-GFP and ALBA5-GFP localization in leaf mesophyll cells**. GFP fluorescence (green) and chlorophyll fluorescence (red) in leaf mesophyll cells. Scale bars = 20 μm.

**Figure S5. Germination kinetics of Col-0, *alba123-1* and *alba123-2***. Germination (radicle emergence) was scored for Col-0 (*n* = 142), *alba123-1* (n = 173) and *alba123-2* (n = 155), distributed over three different plates for each genotype.

**Figure S6. An independent replicate of rosette growth and flowering-time of Col-0, *alba123-1* and *alba123-2***. Col-0 (n = 29), *alba123-1* (n = 28) and *alba123-2* (n = 29) were grown side-by-side as shown in Figure 7C. **(A)** The rosette area of the three genotypes from 16-to 26-days old. **(B)** The flowering-time of the three genotypes. (**** denotes p<0.0001, “ns” indicates “no significant difference”, Student’s t-test).

**Figure S7. Rosette growth and flowering-time of Col-0, *alba123-1* and *alba123-2* under short-day condition**. Col-0 (n = 29), *alba123-1* (n = 19) and *alba123-2*, (n = 20) were grown under identical conditions to the long-day condition experiment except that the photoperiod was only 10 hours per day. **(A)** Rosette area from 20-to 34-days old plants. The area of Col-0 increased significantly faster than that of *alba123-1* and *alba123-2* (*p<*0.001, linear mixed model, ANOVA and Tukey’s HDS). **(B)** The flowering-time of *Col-0, alba123-1* and *alba123-2* under short-day conditions. The plant was scored as flowering when the bolting shoot reached 1 cm. *** indicates *p*<0.001, “ns” indicates no significance (Student’s t-test).

**Figure S8. The different alternative splicing events and their percentages in Col-0 and *alba123***. The x-axis denotes the samples, the y-axis denotes the percentage. The colors denote the particular splicing event (MXE denotes the Mutually exclusive exons, AS5S denotes the Alternative 5’ Splicing Site, RI denotes the Retained Intron, SE denotes Skipped Exon and A3SS denotes Alternative 3’ Splicing Site.

**Figure S9. The GO enrichment analysis and KEGG pathway enrichment analysis. (A)** The GO enrichment with biological process ontology. The Y-axis is the number of DEGs, the X-axis is the annotated GO terms. **(B)** The KEGG pathway enrichment analysis. The Y-axis is the number of DEGs, the X-axis is the annotated pathways. The pathways related to metabolism were outlined in red.

## References

Alonso JM, Stepanova AN, Leisse TJ, et al. 2003. Genome-wide insertional mutagenesis of Arabidopsis thaliana. Science 301, 653–657.

Alvarez JP, Furumizu C, Efroni I, Eshed Y, Bowman JL. 2016. Active suppression of a leaf meristem orchestrates determinate leaf growth. Elife 5, e15023.

Bell SD, Botting CH, Wardleworth BN, Jackson SP, White MF. 2002. The interaction of Alba, a conserved archaeal chromatin protein, with Sir2 and its regulation by acetylation. Science 296, 148–151.

Boyes DC, Zayed AM, Ascenzi R, Mccaskill AJ, Hoffman NE, Davis KR, Gorlach J. 2001. Growth stage-based phenotypic analysis of Arabidopsis: a model for high throughput functional genomics in plants. The Plant Cell 13, 1499–1510.

Chan CW, Kiesel BR, Mondragon, A. 2018. Crystal Structure of Human Rpp20/Rpp25 Reveals Quaternary Level Adaptation of the Alba Scaffold as Structural Basis for Single-stranded RNA Binding. Journal of Molecular Biology 430, 1403–1416.

Chazotte B. 2011. Labeling nuclear DNA using DAPI. Cold Spring Harb Protocols 2011, pdb prot5556.

Chene A, Vembar SS, Riviere L, Lopez-Rubio J J, Claes A, Siegel TN, Sakamoto H, Scheidig-Benatar C, Hernandez-Rivas R, Scherf A. 2012. PfAlbas constitute a new eukaryotic DNA/RNA-binding protein family in malaria parasites. Nucleic Acids Resarch 40, 3066–3077.

Cho H, Cho HS, Hwang I. 2019. Emerging roles of RNA-binding proteins in plant development. Current Opinion in Plant Biology 51, 51–57.

Clough SJ, Bent AF. 1998. Floral dip: a simplified method for Agrobacterium-mediated transformation of Arabidopsis thaliana. The Plant Journal 16, 735–743.

Curtis MD, Grossniklaus U. 2003. A gateway cloning vector set for high-throughput functional analysis of genes in planta. Plant Physiology 133, 462–469.

Da Costa KS, Galucio JMP, Leonardo ES, Cardoso G, Leal E, Conde G, Lameira J. 2017. Structural and evolutionary analysis of Leishmania Alba proteins. Molecular and Biochemical Parasitology 217, 23–31.

Dupe A, Dumas C, Papadopoulou B. 2014. An Alba-domain protein contributes to the stage-regulated stability of amastin transcripts in Leishmania. Molecular Microbiology 91, 548–561.

Dupe A, Dumas C, Papadopoulou B. 2015. Differential Subcellular Localization of Leishmania Alba-Domain Proteins throughout the Parasite Development. PLoS One 10, e0137243.

Edwards K, Johnstone C, Thompson C. 1991. A simple and rapid method for the preparation of plant genomic DNA for PCR analysis. Nucleic Acids Research 19, 1349

Goodstein DM, Shu S, Howson R, et al. 2012. Phytozome: a comparative platform for green plant genomics. Nucleic Acids Research 40, D1178–1186

Gosai SJ, Foley SW, Wang D, Silverman IM, Selamoglu N, Nelson AD, Beilstein MA, Daldal F, Deal RB, Gregory BD. 2015. Global analysis of the RNA-protein interaction and RNA secondary structure landscapes of the Arabidopsis nucleus. Molecular Cell 57, 376–388.

Goyal M, Banerjee C, Nag S, Bandyopadhyay U. 2016. The Alba protein family: Structure and function. Biochimica et Biophysica Acta 1864, 570–583.

Grillet L, Lan P, Li W, Mokkapati G, Schmidt W. 2018. IRON MAN is a ubiquitous family of peptides that control iron transport in plants. Nature Plants 4, 953–963.

Guo L, Ding J, Guo R, Hou Y, Wang D-C, Huang L. 2014. Biochemical and structural insights into RNA binding by Ssh10b, a member of the highly conserved Sac10b protein family in Archaea. The Journal of Biological Chemistry 289, 1478–1490.

Guo R, Xue H, Huang L. 2003. Ssh10b, a conserved thermophilic archaeal protein, binds RNA in vivo. Molecular Microbiology 50, 1605–1615.

Hands-Taylor KL, Martino L, Tata R, Babon JJ, Bui TT, Drake AF, Beavil RL, Pruijn GJ, Brown PR, Conte MR. 2010. Heterodimerization of the human RNase P/MRP subunits Rpp20 and Rpp25 is a prerequisite for interaction with the P3 arm of RNase MRP RNA. Nucleic Acids Research 38, 4052–4066.

Hentze MW, Castello A, Schwarzl T, Preiss T. 2018. A brave new world of RNA-binding proteins. Nature Reviews Molecular Cell Biology 19, 327–341.

Honkanen S, Jones VAS, Morieri G, Champion C, Hetherington AJ, Kelly S, Proust H, Saint-Marcoux D, Prescott H, Dolan L. 2016. The Mechanism Forming the Cell Surface of Tip-Growing Rooting Cells Is Conserved among Land Plants. Current Biology 26, 3238–3244.

Hothorn T, Bretz F, Westfall P. 2008. Simultaneous inference in general parametric models. Biometrical Journal 50, 346–363.

Jelinska C, Conroy MJ, Craven CJ, Hounslow AM, Bullough PA, Waltho JP, Taylor GL, White MF. 2005. Obligate heterodimerization of the archaeal Alba2 protein with Alba1 provides a mechanism for control of DNA packaging. Structure 13, 963–971.

Laurens N, Driessen RPC, Heller I, Vorselen D, Noom MC, Hol FJH, White MF, Dame RT, Wuite GJL. 2012. Alba shapes the archaeal genome using a delicate balance of bridging and stiffening the DNA. Nature Communications 3, 1328.

Lewandowska M, Wawrzynska A, Moniuszko G, et al. 2010. A contribution to identification of novel regulators of plant response to sulfur deficiency: characteristics of a tobacco gene UP9C, its protein product and the effects of UP9C silencing. Molecular Plant 3, 347–360.

Madej T, Lanczycki C J, Zhang D, Thiessen PA, Geer RC, Marchler-Bauer A, Bryant SH. 2014. MMDB and VAST+: tracking structural similarities between macromolecular complexes. Nucleic Acids Research 42, D297–303.

Mani J, Guttinger A, Schimanski B, Heller M, Acosta-Serrano A, Pescher P, Spath G, Roditi I. 2011. Alba-domain proteins of Trypanosoma brucei are cytoplasmic RNA-binding proteins that interact with the translation machinery. PLoS One 6, e22463.

Marondedze C, Thomas L, Serrano NL, Lilley KS, Gehring C. 2016. The RNA-binding protein repertoire of Arabidopsis thaliana. Scientific Reports 6, 29766.

Ozdilek BA, Thompson VF, Ahmed NS, White CI, Batey RT, Schwartz JC. 2017. Intrinsically disordered RGG/RG domains mediate degenerate specificity in RNA binding. Nucleic Acids Research 45, 7984–7996.

Palm D, Simm S, Darm K, Weis BL, Ruprecht M, Schleiff E, Scharf C. 2016. Proteome distribution between nucleoplasm and nucleolus and its relation to ribosome biogenesis in Arabidopsis thaliana. RNA Biology 13, 441–454.

Patterson K, Walters LA, Cooper AM, Olvera JG, Rosas MA, Rasmusson AG, Escobar MA. 2016. Nitrate-Regulated Glutaredoxins Control Arabidopsis Primary Root Growth. Plant Physiology 170, 989–999.

Perez-Diaz L, Silva TC, Teixeira SMR. 2017. Involvement of an RNA binding protein containing Alba domain in the stage-specific regulation of beta-amastin expression in Trypanosoma cruzi. Molecular and Biochemical Parasitology 211, 1–8.

Reichel M, Liao Y, Rettel M, Ragan C, Evers M, Alleaume AM, Horos R, Hentze MW, Preiss T, Millar AA. 2016. In Planta Determination of the mRNA-Binding Proteome of Arabidopsis Etiolated Seedlings. The Plant Cell 28, 2435–2452.

Reiner R, Alfiya-Mor N, Berrebi-Demma M, Wesolowski D, Altman S, Jarrous N. 2011. RNA binding properties of conserved protein subunits of human RNase P. Nucleic Acids Research 39, 5704–5714.

Silverman IM, Li F, Gregory BD. 2013. Genomic era analyses of RNA secondary structure and RNA-binding proteins reveal their significance to post-transcriptional regulation in plants. Plant Science 205-206, 55–62.

Steffen A, Elgner M, Staiger D. 2019. Regulation of flowering time by the RNA-binding proteins AtGRP7 And AtGRP8. Plant Cell Physiol. Jun 26. pii: pcz124.

Subota I, Rotureau B, Blisnick T, Ngwabyt S, Durand-Dubief M, Engstler M, Bastin P. 2011. ALBA proteins are stage regulated during trypanosome development in the tsetse fly and participate in differentiation. Molecular Biology of the Cell 22, 4205–4219.

Szostak E, Gebauer F. 2013. Translational control by 3’-UTR-binding proteins. Briefings in Functional Genomics 12, 58–65.

Tanaka T, Padavattan S, Kumarevel T. 2012. Crystal structure of archaeal chromatin protein Alba2-double-stranded DNA complex from Aeropyrum pernix K1. The Journal of Biological Chemistry 287, 10394–10402.

Torti S, Fornara F, Vincent C, Andres F, Nordstrom K, Gobel U, Knoll D, Schoof H, Coupland G. 2012. Analysis of the Arabidopsis shoot meristem transcriptome during floral transition identifies distinct regulatory patterns and a leucine-rich repeat protein that promotes flowering. The Plant Cell 24, 444–462.

Vasilyev N, Polonskaia A, Darnell JC, Darnell RB, Patel DJ, Serganov A. 2015. Crystal structure reveals specific recognition of a G-quadruplex RNA by a beta-turn in the RGG motif of FMRP. Proceedings of the National Academy of Science USA 112, E5391–E5400.

Verma JK, Gayali S, Dass S, Kumar A, Parveen S, Chakraborty S, Chakraborty N. 2014. *OsAlba1*, a dehydration-responsive nuclear protein of rice (*Oryza sativa* L. ssp. indica), participates in stress adaptation. Phytochemistry 100, 16–25.

Verma JK, Wardhan V, Singh D, Chakraborty S, Chakraborty N. 2018. Genome-wide identification of the Alba gene family in plants and stress-responsive expression of the rice Alba genes. Genes (Basel) 9, 183

Wardleworth BN, Russell RJ, Bell SD, Taylor GL, White MF. 2002. Structure of Alba: an archaeal chromatin protein modulated by acetylation. EMBO Journal 21, 4654–4662.

Wickham H. 2016. ggplot2: Elegant Graphics for Data Analysis, New York, Springer-Verlag.

Wu J, Niu S, Tan M, et al. 2018. Cryo-EM Structure of the Human Ribonuclease P Holoenzyme. Cell 175, 1393–1404 e11.

Wu Z, Zhu D, Lin X, et al. 2016. RNA Binding Proteins RZ-1B and RZ-1C Play Critical Roles in Regulating Pre-mRNA Splicing and Gene Expression during Development in Arabidopsis. The Plant Cell 28, 55–73.

Yuan W, Zhou J, Tong J, Zhuo W, Wang L, Li Y, Sun Q, Qian W. 2019. ALBA protein complex reads genic R-loops to maintain genome stability in *Arabidopsis*. Scientific Advances 5, eaav9040.

Zhang Z, Boonen K, Ferrari P, Schoofs L, Janssens E, van Noort V, Rolland F, Geuten K. 2016 UV crosslinked mRNA-binding proteins captured from leaf mesophyll protoplasts. Plant Methods 12, 42.

Zuo Y, Feng F, Qi W, Song R. 2019. Dek42 encodes an RNA binding protein that affects alternative pre-mRNA splicing and maize kernel development. Journal of Integrative Plant Biology 61, 728–748.

